# The tumor suppressor p53 promotes carcinoma invasion and collective cellular migration

**DOI:** 10.1101/380600

**Authors:** Shijie He, Christopher V. Carman, Jung Hyun Lee, Bo Lan, Stephan Koehler, Lior Atia, Chan Young Park, Jae Hun Kim, Jennifer A. Mitchel, Jin-Ah Park, James P. Butler, Sam W. Lee, Jeffrey J. Fredberg

**Affiliations:** Harvard T.H. Chan School of Public Health, 665 Huntington Avenue, Boston, Massachusetts 02115, USA; Massachusetts General Hospital and Harvard Medical School, Building 149, 13th Street, Charlestown, Massachusetts 02129, USA

**Author notes:** These authors contribute equally to this work (Shijie He, Christopher V. Carman).

## Abstract

**Summary:** Loss of function of the tumor suppressor p53 is generally thought to increase cell motility and invasiveness. Using 2-D confluent and 3-D spheroidal cell motility assays with bladder carcinoma cells and colorectal carcinoma cells, we report, to the contrary, that loss of p53 can decrease cell motility and invasion.

**Abstract:** For migration of the single cell studied in isolation, loss of function of the tumor suppressor p53 is thought to increase cell motility. Here by contrast we used the 2-D confluent cell layer and the 3-D multicellular spheroid to investigate how p53 impacts dissemination and invasion of cellular collectives. We used two human carcinoma cell lines, the bladder carcinoma EJ and the colorectal carcinoma HCT116. We began by replicating single cell invasion in the traditional Boyden chamber assay, and found that the number of invading cells increased with loss of p53, as expected. In the confluent 2-D cell layer, however, for both EJ and HCT, speeds and effective diffusion coefficients for the p53 null types compared to their p53 expressing counterparts were significantly smaller. Compared to p53 expressers, p53 null cells exhibited more organized cortical actin rings together with reduced front-rear cell polarity. Furthermore, loss of p53 caused cells to exert smaller traction forces upon their substrates, and reduced formation of cryptic lamellipodia. In a 3-D collagen matrix, p53 consistently promoted invasion of the multicellular spheroids into surrounding matrix. Together, these results show that p53 expression in these carcinoma model systems increases collective cellular migration and invasion. As such, these studies point to paradoxical contributions of p53 in single cell versus collective cellular migration.

## Introduction

Among human cancers, the tumor suppressor p53 is the most mutated gene and serves not only as an inducer of cancer cell senescence and apoptosis [1,2], but also as a central suppressor of cancer cell migration and metastasis [3–6]. For example, in 3-dimensional (3D) Matrigel assays, loss of p53 increases single cell invasion by enhancing cell contractility [7–10]. In wound healing assays, p53 can decrease the migration distance of leading cells by the inhibition of epithelial-mesenchymal transition (EMT) [11]. In addition, p53 can inhibit cancer cell metastasis by suppressing focal adhesion kinase (FAK) [12] and preventing degradation of the extracellular cell matrix (ECM) [3,13].

Most studies to date have emphasized effects of p53 on single cell invasion in the Matrigel-coated Boyden chamber assay [7–10]. It is now recognized, however, that metastatic disease is dominated by collective cellular migration rather than single cell migration [14–16]. In the case of collective cellular migration, the cell-cell interactions can be quite strong and highly cooperative [17–21]. Moreover, the cellular collective can become jammed, immobile, and solid-like, or unjammed, mobile, and fluid-like [18,22,23]. In the case of single cell migration, by contrast, none of these potent mechanisms are operative. It remains unclear, however, how p53 functions in the context of such collective phenomena.

To address that issue, here we studied migration and invasion in 2D confluent cell layers and 3D multicellular spheroids. Two human cell lines were used, the bladder carcinoma EJ and the colorectal carcinoma HCT116. We first replicated single cell invasion assays in the Boyden chamber and found results consistent with previous studies [7–9]; loss of p53 increased the invasion of the single carcinoma cell. To our surprise, however, loss of p53 in either EJ or HCT 116 cells suppressed cellular dissemination in 2-D confluent cell layers. In that case, loss of p53 was associated with reduced lamellipodia formation and weaker cell-substrate interactions. To better mimic tumor biology we also conducted studies using 3D multicellular spheroids embedded in collagen matrix. We found these results in the 3D multicellular spheroidal assay to be consistent with the 2-D confluent assay. These results, taken together, demonstrate paradoxical contributions of p53 in single cell versus collective cell migration.

## Results

### In 2D confluent cell layers, p53 increases collective cellular motility

To determine the function of p53 in collective cell motility, we used both gain and loss of p53 function in colorectal and bladder carcinoma cell lines: stable wild type (p53^+/+^) and stable p53 null (p53^−/−^) HCT 116 and Tet-off inducible EJ cell line (Methods). In EJ cell line, p53 knocking out (EJ p53 off) was established by the addition of doxycycline to the culture media. We began by replicating assays of single cell invasion in the Matrigel-coated Boyden chamber as reported in previous studies [7–10], and found consistent results; loss of p53 increased cell invasion (125±53 versus 415±101 cells per well for EJ, p = 0.009 249±65 versus 891±239 for HCT 116, p = 0.03) (Fig. S1).

We then went on to assays of cell migration in the 2-D confluent cell layer. The confluent cell layer was cultured on 1.2 kPa polyacrylamide gel, and red fluorescent beads embedded in the gel surface. We used Leica DMI8 with living cell culture system to require phase images of the cell layer and fluorescent images of the read beads at 10 minutes interval for 24 h. Based on the phase images we calculated cell velocity and displacement in the confluent layer by using optical flow [24]. Using Traction Force Microscopy, we measured the traction forces exerted by the confluent layer on the gel (Methods and Fig. 1A). To our surprise, mean cell speeds for the p53 null version (EJ p53 off and HCT116 p53^−/−^) were significantly smaller compared to their p53 expressing counterparts (0.12±0.003 versus 0.16±0.01μm min^−1^, p=0.002 for EJ; 0.11±0.02 versus 0.18±0.02 m min^−1^, p=0.0003 for HCT 116; Figs. 1B and 1C). When cell motions were expressed as an effective diffusion coefficient, the diffusivities of p53 null cells were consistently smaller than those of p53 expressing controls (0.36±0.07 versus 0.77±0.23, μm^2^ min^−1^, p=0.003 for EJ; 0.20±0.05 versus 0.43±0.06 μm^2^ min^−1^, p=0.0003 for HCT 116 in Figs. 1D and 1E, Movies S1 and S2).

**Fig 1.**
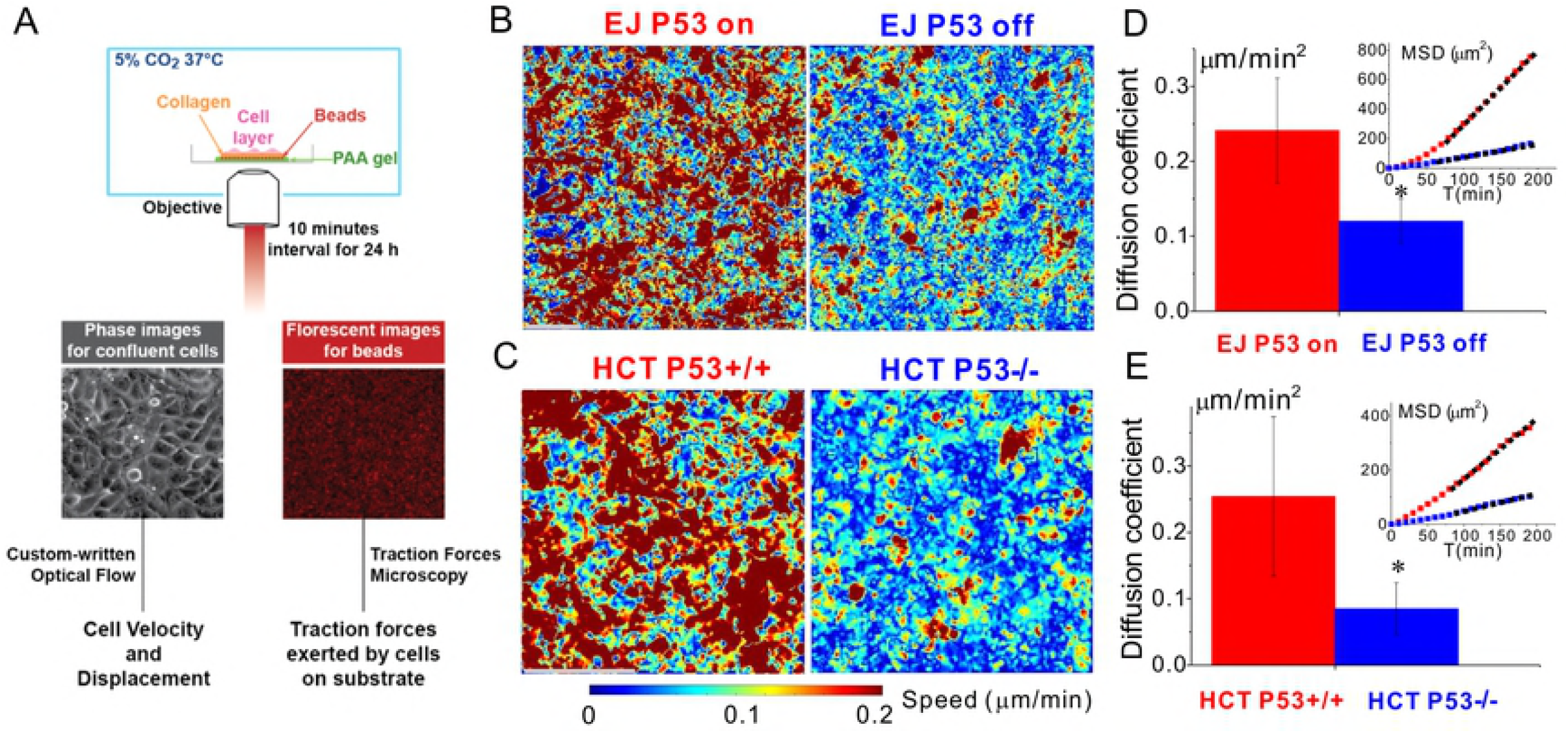
Loss of p53 reduces carcinoma motility and diffusion in confluent 2D assay system. **A**. Cell velocity and displacement in the confluent cell layer were calculated based on the cell phase images by using custom-written optical flow. Traction forces exerted by the confluent cell layer on the Polyacrylamide (PAA) gel were derived from the florescent images of beads by using traction force microscopy (Method). (**B** and **C**) Speed maps shows that loss of p53 decreases the speed of EJ cells and HCT116 cells. During 24h, their mean speeds are 0.12±0.003 versus 0.16±0.008 μm min^−1^, p=0.002 for EJ; 0.11±0.018 versus 0.18±0.025 μm min^−1^, p=0.0003 for HCT 116. (**D** and **E**) The diffusion coefficients, D, of EJ p53 off cells (HCT 116 p53^−/−^ cells) were smaller than EJ p53 on cells (HCT 116 p53^+/+^ cells). As such, p53 null cells were less diffusive than p53 expressing counterparts. D was calculated by linear fitting the mean squared displacements MSD= 4D*T+b after 70 min, as shown in the representative MSD in the insets. Sample number n=6~8, * represents p<0.05, Scale bar, 100μm.

### P53 null cells show weaker cell-substrate interactions

Cell-substrate interaction plays a critical role in collective cellular migration. To our knowledge, the effects of p53 on cell-substrate interactions have not yet been reported. Over 24h we continuously measured the traction forces. As shown in the representative traction maps (Figs. 2A and 2B) the p53 null cells had fewer hot spots than did p53 expressing cells. Root mean square traction (RMST) from p53 null cells was smaller than in p53 expressing counterparts (Figs. 2C and 2D). Mean RMST was 23.3±0.3 versus 35.9±4.1 Pa, p=0.01 for EJ; 8.2±0.3 versus 10.2±0.3 Pa, p=0.1 for HCT 116. Thus expressing p53 caused cells to exert larger traction forces upon their substrates.

**Fig 2.**
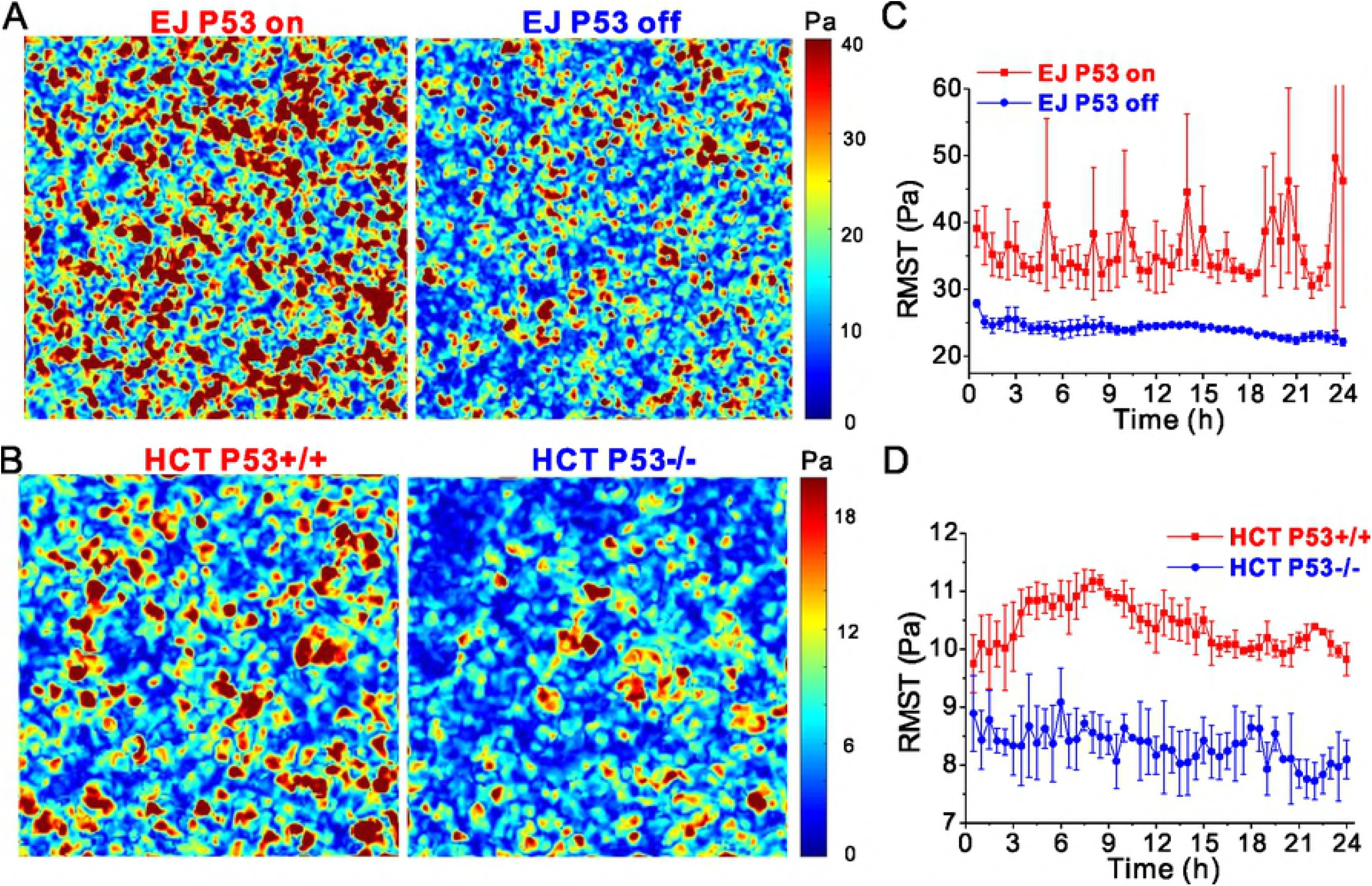
P53 increases traction exerted by the cell layer on the substrate. (**A** and **B**) Representative traction maps from the p53 null cells have fewer hot spots than do p53 expressing cells. (**C** and **D**) Root mean squared tractions (RMST) were continuously measured during 24h. Expressing p53 increases RMST, and mean RMST are 23.3±0.3 Pa versus 35.8±4.1 Pa, p=0.01 for EJ cells, and 8.2±0.3 versus 10.2±0.3 Pa, p=0.1 for HCT 116 cells.

### P53 null cells exhibit more organized cortical actin rings together with reduced front-rear cell polarity

In the bladder carcinoma EJ p53 on and EJ p53 off, and in the colorectal carcinoma HCT116 p53^+/+^ and HCT116 p53^−/−^, western blots confirmed expected p53 expression or lack thereof (Fig. 3A and Fig. 4A). Fluorescent staining with phalloidin showed that the apical cortical F-actin rings found in both of the p53 null carcinomas (EJ p53 off and HCT116 p53^−/−^) were more round and intact than their p53 expressing counterparts (Fig. 3B and Fig. 4B). However, expression of E-cadherin varied in EJ and HCT 116. In western blotting and fluorescent staining assays, E-cadherin was not detectably expressed in both EJ p53 on and EJ p53 off (Fig. 3A, fluorescent staining not shown). Alternatively, E-cadherin was expressed and localized to the cell-cell junctions in both HCT116 p53^+/+^ and HCT116 p53^−/−^, with the latter showing ~2.5-fold greater levels of E-cadherin than the former (Fig. 4A and Fig. 4C). While a consistent relationship existed between loss of p53, reduced motility and increased cortical actin in these two different carcinomas, no such correlations was found regarding their E-cadherin expression.

**Fig 3.**
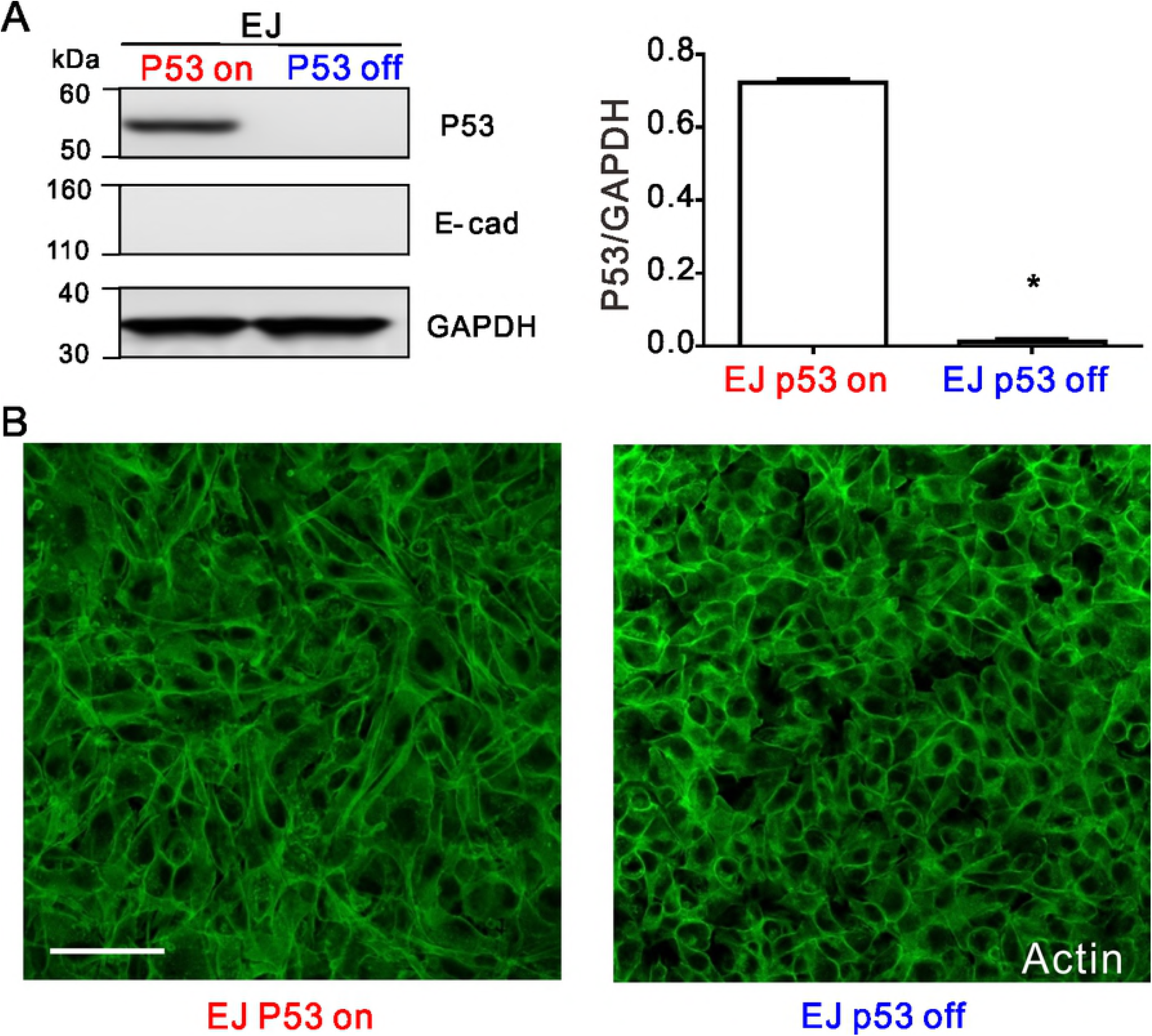
The expression of p53, E-cadherin and actin in the EJ cells. (**A**) Western blot cropped from same gel confirms that the p53 expression in the EJ p53 off cells are null, and E-cad is not detectable. The full-length blots are presented in Fig S2A. (**B**) Fluorescent staining shows that actin rings are more organized in the EJ p53 off cells than in EJ p53 on cells. E-cadherin is also not detectable in fluorescent staining (data not shown). Scale bar, 100μm. The fluorescent images represent at least six field views from two experiments. These western blot and fluorescent staining for both EJ and HCT 116 are performed in the same condition (Methods), and as shown in Fig. 4 E-cadherin expression in HCT 116 cells serves as the positive control.

**Fig 4.**
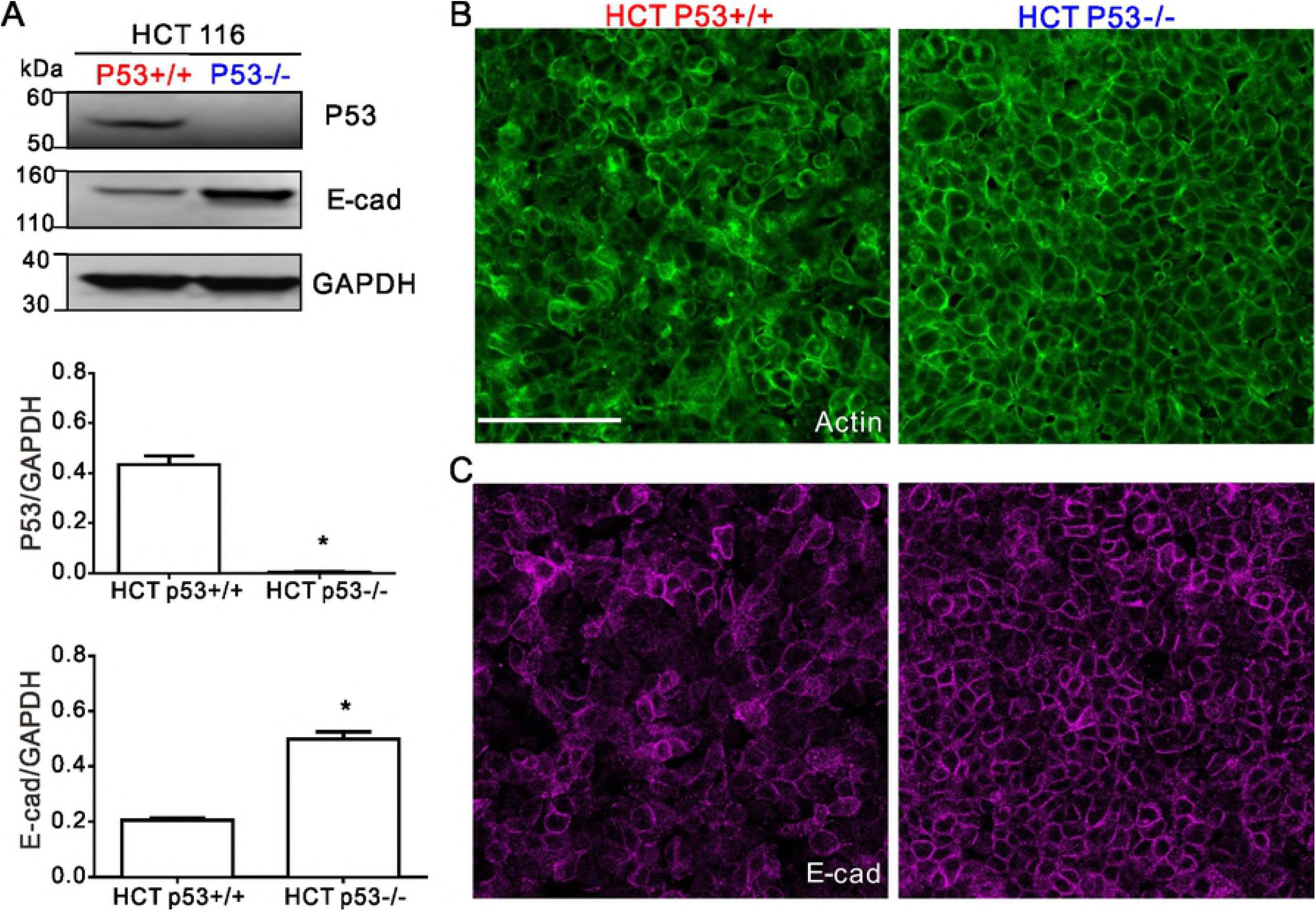
The expression of p53, E-cadherin and actin in the HCT116 cells. (**A**) Western blot cropped from same gel confirms that the p53 expression in HCT 116 p53^−/−^ cells are null. E-cadherin expressions in HCT 116 p53^−/−^ cells are ~2.5-fold higher than HCT 116 p53^+/+^ cells. The full-length blots are presented in Fig S2B. (**B**) Fluorescent staining shows that actin rings are more organized in the p53^−/−^ cells than in the p53^+/+^ cells. (**C**) Fluorescent staining shows that E-cadherin is located at the cell-cell junction in both the p53^+/+^ and p53^−/−^ cells, and more in the p53^−/−^ cells than the p53^+/+^ cells. Scale bar, 100μm.

### P53 null cells show reduced formation of cryptic lamellipodia

Fluorescent images of F-actin were obtained using an inverted confocal microscope (Leica SP8, Method). Near the basal plane of both EJ and HCT 116 carcinomas, the F-actin images showed cryptic lamellipodia (the F-actin tips labeled by asterisks in Figs. 5A and 5B). We found that loss of p53 was associated with reduced appearance of the cryptic lamellipodia, which are typically associated with collective cell migration. Moreover, normalized intensity measurements of fluorescent F-actin showed that near the basal plane expressing p53 increased F-actin intensity, 1.5±0.2 times for EJ cells, 1.8±0.4 times for HCT 116 cells (Figs. 5C and 5D).

**Figure 5.**
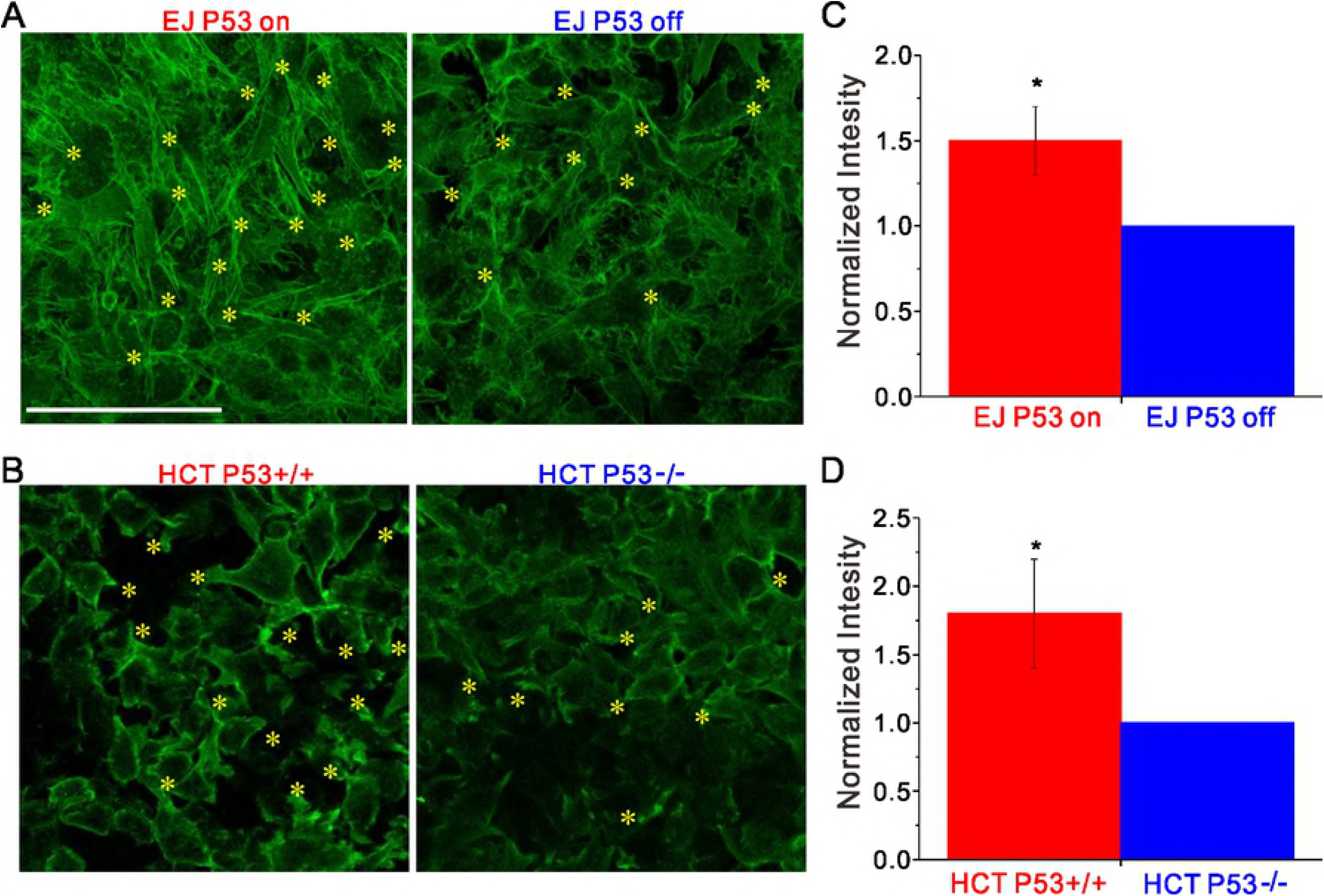
Near basal plane of the cell layers p53 null cell shows reduced appearance of cryptic lamellipodia. (**A** and **B**) Near the basal plane of the cell layer, F-actin tips indicated by the asterisks showed the appearance of cryptic lamellipodia. For both EJ and HCT 116, p53 null cells showed reduced appearance of the cryptic lamellipodia. (**C**) F-actin intensity for EJ p53-on cells are 1.5±0.2 times than that for EJ p53-off cells. (**D**) F-actin intensity for HCT 116 p53^+/+^ cells are 1.8±0.4 times than that for HCT 116 p53^−/−^ cells. Scale bar, 100μm.

### P53-null multicellular spheroids exhibit lower 3-D invasiveness

To better mimic tumor biology, we next conducted studies using 3D multicellular spheroids embedded into collagen matrix. The invasiveness of each carcinoma spheroid was assessed via the invaded area (as enveloped by the yellow dotted line in Fig. 6A) normalized by the initial area of the cross section of the spheroid. This metric showed that loss of p53 reduced invasion of the carcinoma spheroids into surrounding matrix (3.8±1.4 versus 8.4±1.5, p<0.0001 for EJ; 1.2±0.1 versus 2.8±0.8, p<0.0001 for HCT 116 in Fig. 6B). Compared to their p53 expressing counterparts, p53 null carcinomas (EJ p53 off and HCT 116 p53^−/−^) exhibited less frequent escape of individual cells from the spheroids and less efficient dissemination (Movies. S3 and S4). These results from 3D multicellular spheroids are consistent with those from 2D confluent layer, which both suggest that p53 promotes carcinoma invasion and collective cellular migration.

**Fig 6.**
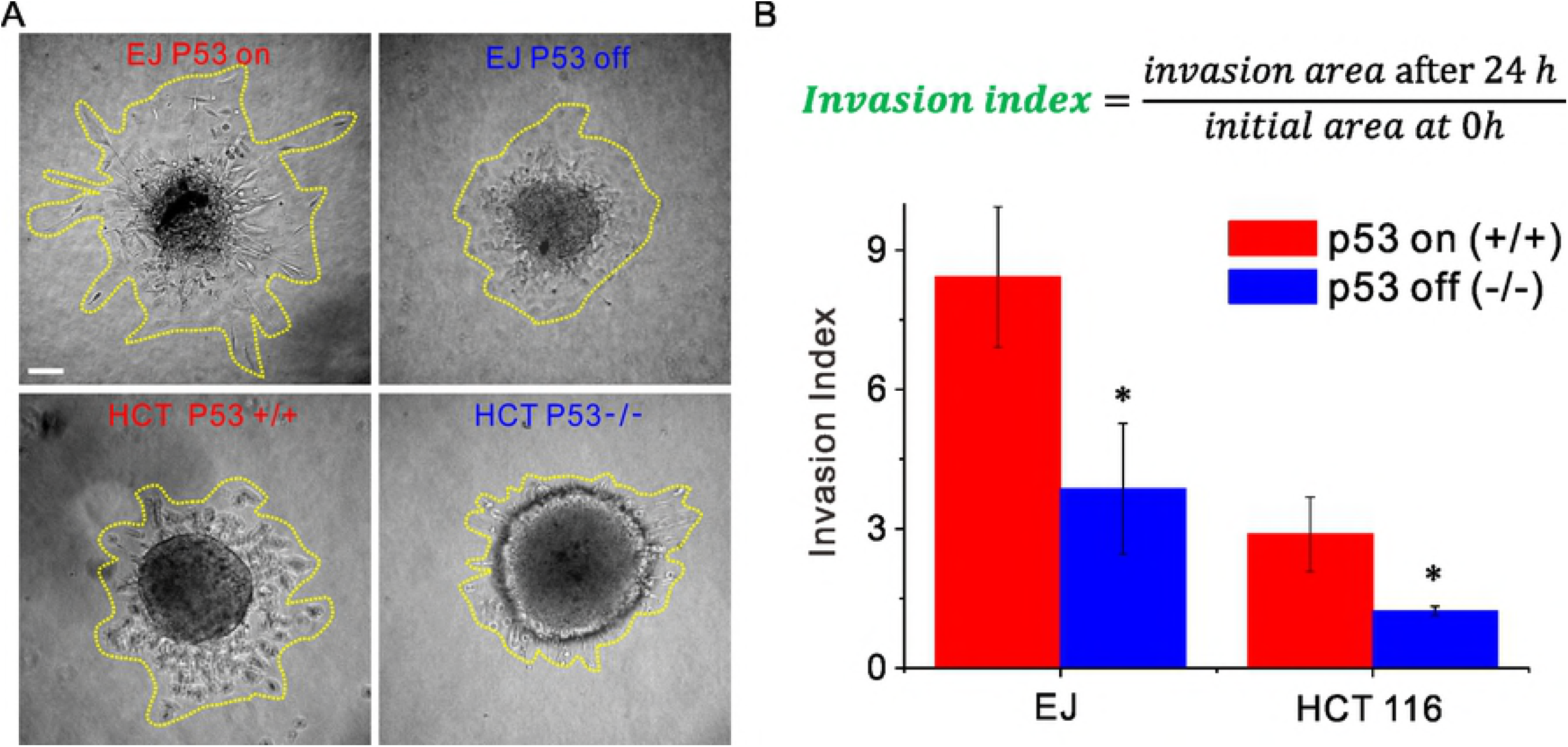
In 3D collagen matrix, the multicellular spheroids formed by the p53 null cells are less invasive than the ones formed by the p53 expressing counterparts. (**A**) Representative images show the invasion of the multicellular spheroids. The p53 null spheroids invade less than the p53 expressing cells in the both cell lines. The yellow lines indicate the area invaded by cells. (**B**) Invasion index is the area invaded by cells after 24h divided by the initial area. This invasion index for p53 null multicellular spheroids is lower than that from p53 expressing cells (3.8±1.4 versus 8.4±1.5, p<0.0001 for EJ; 1.2±0.1 versus 2.8±0.8, p<0.0001 for HCT 116). Collagen density, 1.5 mg/ml. Scale bar, 100μm.

## Discussion

Using bladder and colorectal carcinoma cell lines, here we confirmed that p53 suppresses single cell invasion in the Boyden chamber assay [7–9]. Paradoxically, however, in both the 2D and 3D assays p53 increased collective cellular migration and invasion. Together, these results suggest that the function of p53 in cell motility depends on context, that is, single cell migration versus collective cellular migration.

Function of p53 depends on its target genes and proteins, which can exert paradoxical, context-dependent, effects on the same cellular process, including apoptosis, metabolism, differentiation and migration [25]. For example, evidence for regulation of cell migration by p53 is conflicting. The preponderance of the evidence suggests that the tumor suppressor p53 attenuates cell motility and invasiveness through pathways [3–6] such as suppressing the RhoA-Rock or Rac1 [7,8,10] and inhibiting epithelial-mesenchymal transition (EMT) [11,26]. Limited evidence suggests, to the contrary, that p53 promotes in vitro and in vivo invasion of ovarian carcinoma cells isolated from PTEN; KRas mice [27], and also promotes migration and invasion of human lung, colorectal carcinoma and osteosarcoma cells by activating Rac1 or Rap2a [28,29]. These studies exemplify the paradoxical regulation by p53 in the cell migration from the perspective of different target signal pathways. As regards the contexts of single cell versus collective cellular migration, however, the function of p53 has remained unclear. Here we suggest paradoxical contributions of p53 in single cell versus collective cellular migration.

### P53 promotes the dissemination and invasion of carcinoma cellular collectives

In the cellular collective, results from both bladder and colorectal carcinomas support the notion of the tumor suppressor p53 as a promoter of dissemination and invasion. To manipulate p53, two distinct methods were used in the two human carcinoma cell lines: p53 tet-off system of the bladder carcinoma EJ, and wild-type p53 cells and knock out p53 cells of colorectal carcinoma HCT 116 (Method). To quantify the cell dissemination in the 2-D confluent cell layer, we measured cell speed, mean squared displacement (MSD), and the diffusion coefficient (Fig. 1). To our surprise, in both EJ and HCT 116 these metrics from the cells expressing p53 were all higher than those from the p53 null counterparts. As such, the tumor suppressor p53 was associated with faster collective cellular migration and easier escape of trapped cells from their neighbors (Movies S1 and S2). Both the bladder and colorectal carcinoma spheroids in collagen matrix, which better mimic tumor microenvironment, showed consistent results; the tumor suppressor p53 promotes the carcinoma spheroid to invade a larger area (Fig. 6). These results demonstrate that the tumor suppressor p53 promotes carcinoma cell escape from their neighbors and more efficient invasion into matrix.

### P53 increases formation of cryptic lamellipodia and migratory traction forces

Our results show that p53 promotes the dissemination and invasion of the cellular collectives. Compared to p53 expressing counterparts, the p53 null carcinomas shared the features of having highly organized rings of cortical F-actin, and more rounded and less polarized cell shape (Fig. 3B and Fig. 4B). These results are consistent with the general consensus that loss of p53 promotes cellular rounding [7,30]. Moreover recent studies have established cell rounding (i.e., lower aspect ratio) as a key physical mechanism to make cell collectives less diffusive and more jammed by increasing the energy barrier for cells to escape from their neighbors (i.e., exchange neighbors in confluent layers) [22,31,32].

Cryptic lamellipodia represent a critical structure for collective cellular migration [33]. Near the basal planes of both the bladder and colorectal carcinoma cellular collectives, fluorescent images of F-actin showed that loss of p53 was associated with the reduced formation of the cryptic lamellipodia (Fig. 5A and 5B), which is consistent with the lower motility of the p53 null cells. These results support the notion that p53 can activate the formation of cryptic lamellipodia to promote cell dissemination and invasion in the cellular collectives.

To our knowledge these studies are the first to quantify the mechanical effects of p53 on cell-substrate interaction. Measurements in both EJ and HCT 116 suggest that the loss of p53 consistently causes these carcinoma cells to exert smaller traction forces on their substrate (Fig. 2). It remains unclear, however, if reduction in traction forces might be attributable to reduced lamellipodia formation and less motility.

### Discordant relationship between p53-dependant regulation of collective carcinoma migration and E-cadherin expression

Many studies suggest that p53 can prevent epithelial-mesenchymal transition (EMT) and increase E-cadherin expression to decrease cancer cell motility [11,26,34,35]. Nevertheless, at least one study [9] suggests that loss of p53 does not decrease the E-cadherin expression. Our current studies in the 2-D confluent cell layers also show a discordant relationship between p53 and E-cadherin expression. The bladder carcinomas do not express E-cadherin regardless of expressing p53 or not, which suggests that in this context the effects of p53 on the motility of collective carcinomas are not mechanistically related to E-cadherin expression. Nevertheless, in the bladder carcinoma collective it remains unclear whether the effects of p53 on cell migration are associated with the epithelial-to-mesenchymal transition (EMT), and whether p53 increases expression of other cadherins, such as P- and N-cadherin. For the colorectal carcinoma cells, loss of p53 increases E-cadherin expression, which might contribute to increased cell-cell interaction so as to cage cells by their neighbors, although theoretical models suggest, to the contrary, that increasing cell-cell adhesion decreases the energy barrier for cells to escape from their neighbors [31]. These results suggest that regulation of E-cadherin expression by p53 is context-dependent, and may differ for different carcinoma types.

In conclusion, this study points to paradoxical contributions of the tumor suppressor p53 in single cell versus collective cellular migration. P53 inhibits migration of the single cell studied in isolation, but both in 2D and 3D assays p53 promotes the dissemination and invasion of the confluent cellular collective. P53 was also associated with changes in the F-actin cytoskeleton, cell morphology, lamellipodia, traction forces as well as E-cadherin expression, all of which are thought to be linked with migration of the cellular collective (Fig. S3).

Mechanism by which p53 regulates the collective carcinoma cell migration remains unclear. P53 suppresses RhoA to inhibit the amoeboid migration of single cell [7]. But in the context of collective cellular migration, it remains unclear if p53 suppresses RhoA [7], or activates Rac1 [28] to cause elongation, formation of cryptic lamellipodia [36], and stronger cell-substrate interactions, all of which might promote migration of the cellular collectives. In addition unanswered questions concern context-dependent regulation by p53 in senescence and apoptosis. P53 mutations show gain of function in promoting cancer cell invasion [4], but how these p53 mutations function in cellular collectives remains unclear. Evidence presented here demonstrates that the tumor suppressor p53 paradoxically promotes carcinoma invasion and collective cell migration, and, in the case of cancer therapeutics, implies the possibility that p53 might act to promote collective carcinoma migration rather than suppress it.

## Methods

### Cell culture

Human bladder carcinoma EJ cells were cultured in Dulbecco’s modified Eagle’s medium (DMEM, Corning, 10-013-CV) containing 10% Fetal Bovine Serum (FBS, Atlanta Biologics), 100 μg/ml of Hygromycin (Sigma, H3274), and 200 μg/ml Geneticin G418 (Teknova, G5005). P53 knock-out EJ cells (EJ p53 off) and p53 expressing EJ cells (EJ p53 on) were established respectively by the presence and the absence of 1 μg/ml doxycycline (Sigma, D9891) in the culture media. Human colorectal carcinoma HCT116 cells were cultured in DMEM containing 10% FBS. Both EJ and HCT 116 cells were maintained at 5% CO2 and 37°C. All the human cell lines and experimental protocols were approved by Harvard Institute Review Board and carried out in accordance with the relevant guidelines and regulations. The datasets generated during the current study are available from the corresponding author on reasonable request.

### 2D confluent cell layer assay

Polyacrylamide gel substrates (Young’s modulus, 1.2kPa, thickness, 100μm) were fabricated on dishes, and for the traction measurement 0.5μm fluorescent red beads (Invitrogen, F8823) were embedded near the gel surface [21]. On the gel we then coated an 8×8mm^2^ square region with 0.1 mg/ml collagen (Advanced Biomatrix, 5005). 80μl cell suspension (5×10^5^ cells/ml for EJ, and 8×10^5^ cells/ml for HCT 116) were seeded on the collagen coated region. After 24h, both phase contrast images and fluorescent images were captured for the cells and the beads respectively via an inverted fluorescent microscope (Leica DMI8) (10 min interval for 48 h at 5% CO2 and 37°C). The phase images were used to quantify cell motion via our custom-written software based on the function in MATLAB termed as opticalFlowFarneback, and we reported the cell velocity map and the mean squared displacement (MSD) as shown in Fig. 1. The fluorescent images of the red beads were used to calculate the gel deformation, as described in following.

### Traction Force Microscopy

Traction was quantified by traction force microscopy (TFM) [37]. At the end of the 2D confluent cell layer assay, cells were detached with trypsin (Corning, 25-052-CI), then we captured the fluorescent images of the red beads as the reference for the gel deformation. The gel deformation was calculated via our custom-written particle image velocimetry software. Based on the gel deformation field, the traction was computed via constrained Fourier transform traction [37]. We reported the traction map and root mean squared traction (RMST) in Figs. 4 and 5.

### 3D multicellular spheroid assay

A 200μl cell suspension (2×10^5^ cells/ml for EJ p53 on, 5×10^5^ cells/ml for HCT 116) was cultured in an ultra-low attachment 96-well plate (VWR, 29443-034) to form the multicellular spheroid. After 48h, each spheroid was carefully pipetted into 1.5mg/ml collagen matrix (Advanced Biomatrix, 5005). Cell culture medium was used to adjust the collagen concentration, and the collagen matrix was equilibrated through 10X PBS (volume ratio, 1:10 between 10X PBS and collagen) and 1M NaOH (0.5% of total matrix volume). These processes were performed on ice to avoid collagen polymerization. We then moved the collagen matrix into 37°C incubator to induce collagen polymerization. After 1h, we monitored the invasion of the spheroids via Leica microscope (Leica DMI8).

### Immunofluorescence and confocal laser-scanning microscopy

For immunofluorescence analysis of cell lines, cell layers from the 2-D assays were fixed in 4% paraformaldehyde/PBS for 10 min, permeabilized and blocked in 0.2% Triton X-100/PBS containing milk for 20 min. The cell layers were stained with primary E-cadherin antibody (Invitrogen, 334000) [11], then stained with secondary antibodies Alexa Fluor 488 (Invitrogen, A-11029). F-actin was stained with Phalloidin conjugated with Alexa Flour 594 (Invitrogen, A12381). LSM (laser scanning microscopy) images were captured by using inverted confocal microscope (Leica SP8, 40x/0.8 oil objective). These images were then processed by using same setting in Image J. All the florescent images represented at least six field views from two experiments.

### Western blot

Protein expression was determined by western blot. All experiments were performed at 72 hours to ensure adequate protein expression. Cells were washed twice with cold PBS and then cell lysates were collected on ice with 10μg phosphatase inhibitor cocktail (Roche). Equal amount of protein lysates from each condition were separated by using NuPAGE 15 well 4-12% Bis-Tris protein gel (Thermo Fisher, MA), then transferred onto nitrocellulose membrane. The membrane was cut into three and probed separately with E-cad antibody (Invitrogen, 334000), p53 antibody (cell signaling, 2527S) and GAPDH antibody (GeneTex, 627408), followed by secondary antibody (Abcam, 205718 & 97040). Here we used GAPDH as our loading control, because the variation of GAPDH expression among different donors/transfections was less than 1 fold (data not shown here). These antibodies have been confirmed by manufacture and previous studies [11,38].

### Single cell invasion assay

The 24-well Boyden chamber (Corning, 354480) contains an 8μm pore size PET membrane which has been coated by Matrigel. We added warmed serum-free culture medium into the interiors of the chambers and the bottoms of the wells, and allowed the system to rehydrate for 2 hours in humidified tissue culture incubator. After rehydration, we carefully removed the medium without disturbing the layer of Matrigel. We prepared the serum-free cell suspension (5×10^4^ cells/ml for EJ, 2×10^5^ cells/ml for HCT 116), and then added 0.5ml into the chamber. We used sterile forceps to transfer the chambers to the wells containing 0.75ml culture medium containing serum as chemoattractant. Cells were incubated in the chambers for 22 hours in 37°C, 5% CO_2_ incubator. After the incubation, we used cotton tipped swabs to scrub the surface of the chamber twice to remove the non-invading cells from the upper surface. Cells were then fixed and stained F-actin and nucleus via the above methods. We counted the nuclei of the entire well bottom through the particle analysis in Image-J. N=4 for each type from two experiments.

## Supporting information

**S1 Fig.**
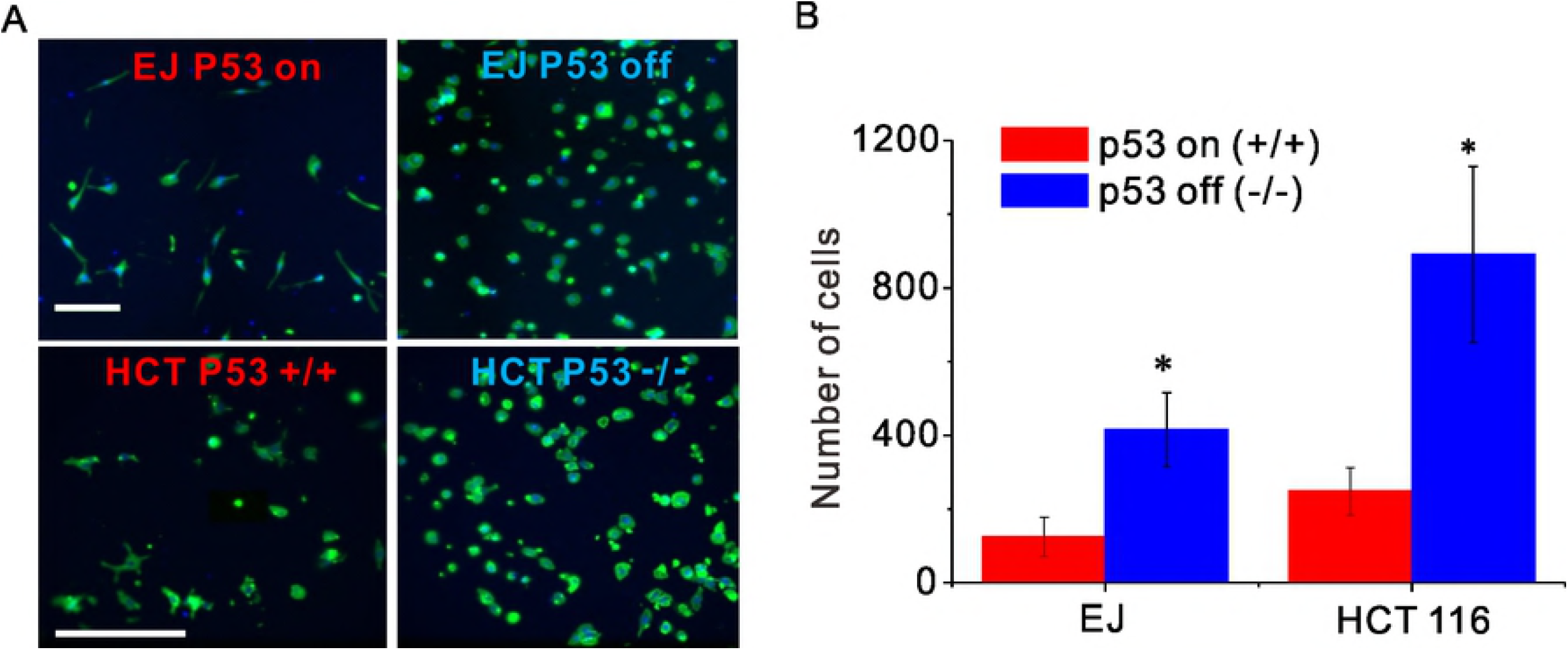
P53 inhibits invasion of the single carcinoma cell. (**A**) shows the representative images for the bottom of the Boyden chamber (green for F-actin, blue for nucleus). The p53 null cells invade more than the p53 expressing counterparts. (**B**) The numbers of the cells per well from the p53 null cells are higher than those from p53 expressing counterparts, (125.0±52.7 versus 415.2±100.7 for EJ; 248.8±64.6 versus 891±238.8 for HCT 116). Scale bar, 200μm.

**S2 Fig.**
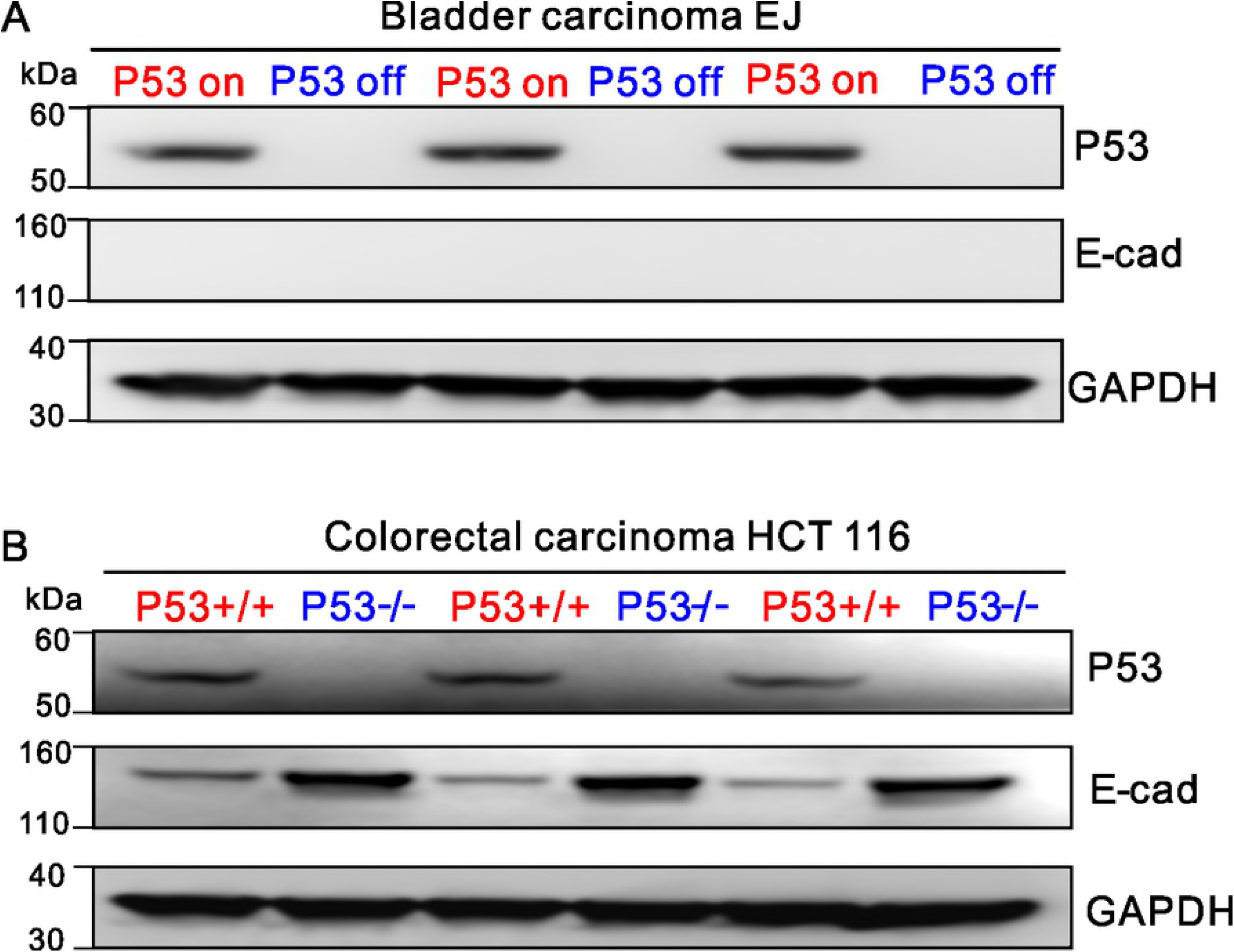
Western blot of p53, E-cadherin and GAPDH for EJ and HCT 116. Exposure time is 10s for GAPDH, 60s for E-cadherin for both EJ and HCT 116 cells, and 10s and 30s for p53 of EJ cells and HCT 116 cells respectively.

**S3 Fig.**
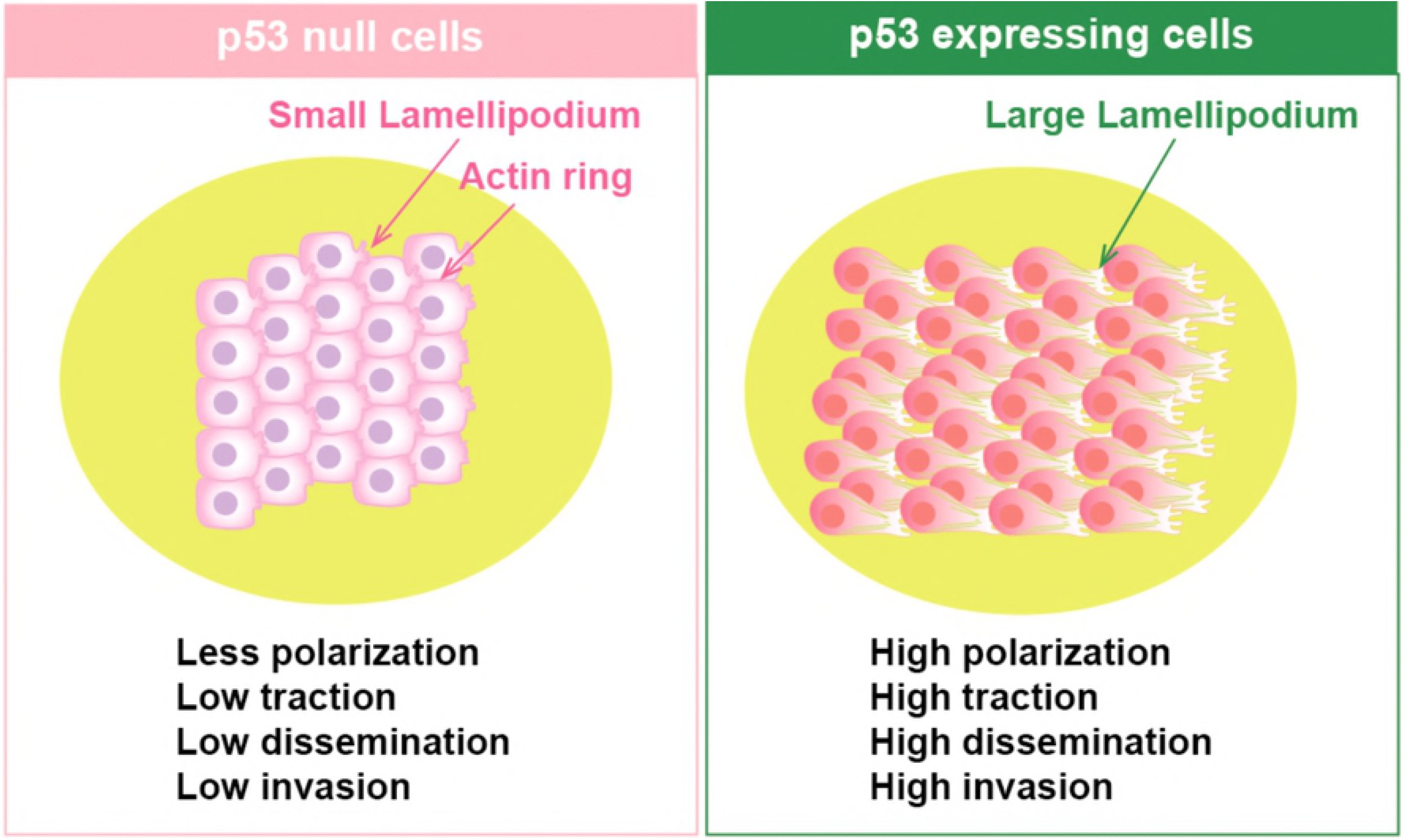
Illustration for the differences between the p53 null and p53 expressing collective cells. Compared to p53 expressers, p53 null cells exhibit more organized cortical actin rings together with reduced front-rear cell polarity and less formation of cryptic lamellipodia. Moreover our study show that p53 increases the traction exerted by the collective cells on substrate, and promotes dissemination and invasion of the collective cells.

**S1 Movie. Cell migration in the 2-D confluent EJ cell layer.**

**S2 Movie. Cell migration in the 2-D confluent HCT 116 cell layer.**

**S3 Movie. Cell invasion of the 3-D EJ spheroid.**

**S4 Movie. Cell invasion of the 3-D HCT 116 spheroid.**

